# A sequence-to-function model to predict T7 transcription rates and redesign T7 expression systems with lowered production of immunogenic RNA byproducts

**DOI:** 10.64898/2026.08.01.742228

**Authors:** James R. McLellan, Howard M. Salis

## Abstract

T7 RNA polymerase is widely used to produce RNA using a canonical T7 promoter; however, it will also bind to low-affinity sites to generate cryptic transcription and produce RNA byproducts, which reduce full-length mRNA purity and yield. When manufacturing therapeutic RNAs for clinical applications, RNA byproducts must be removed using costly downstream purification and can cause adverse immunogenicity. To predict T7 transcription rates and reduce cryptic transcription, we designed 11588 T7 promoters and measured their mRNA levels, spanning a 6300-fold range within *in vitro* transcription reactions. We developed the T7 Promoter Calculator, a sequence-to-function machine learning model that predicts the T7 transcription rate on arbitrary DNA sequence across a 500-fold range with high accuracy (R^2^ = 0.80), accounting for both core and flanking motif sequences. We combined the model with generative design to remove low-affinity T7 sites from a therapeutic T7 expression system, resulting in a 2-fold increase in full-length mRNA purity. The automated design of T7 expression systems to remove undesired RNA byproducts increases mRNA purity and lowers downstream separation costs, while reducing adverse immunogenicity.

## Introduction

RNA therapeutics have emerged as a versatile class of biologics with a wide range of applications across medicine, including the development of protective vaccines and cancer immunotherapies, the treatment of genetic disorders, and genetic modulation to treat cardiovascular and neurological disease^1, 2, 3, 4, 5, 6, 7, 8, 9, 10, 11^. mRNA therapeutics are designed to express proteins *in situ*, for example, to develop adaptive immunity against pathogen or tumor antigens, to express missing enzymes as a replacement therapy, to deliver base editing enzymes to treat disorders, or to express transcription factors for cell reprogramming. As the field advances to routinely translating biologics to the clinic, there is an increasing need to streamline RNA manufacturing and purification processes to reliably create high-purity biologics at low cost.

Therapeutic RNAs are commonly synthesized using T7 RNA polymerase (RNAP) and *in vitro* transcription (IVT) reactions^12, 13^ to produce large quantities of uncapped RNA, using pseudouridine instead of uracil to lower immunogenicity^14^, followed by additional capping reactions^15^. T7 RNAP is widely used because it binds tightly to the canonical T7 promoter and has a fast & highly processive elongation cycle, resulting in large quantities of RNA at low cost^1, 2, 3, 16, 17^. However, T7 IVT will also produce RNA byproducts, including truncated and double stranded RNA (dsRNA), whenever T7 RNAP binds to off-target sites in the DNA template, resulting in cryptic transcription^18, 19, 20^. Long RNA byproducts reduce RNA purity and yield; they are also challenging to separate from full-length mRNAs using liquid chromatography, including affinity chromatography with oligoT resin, because many long RNA byproducts will contain polyA tails^21, 22^. When introduced into the human body, RNA byproducts can have adverse physiological effects, for example, when dsRNAs trigger an antiviral immune response or when truncated mRNAs produce inactive proteins^23, 24, 25^. Notably, the costs of downstream purification are 50-80% of the overall production costs^26^.

This challenge can be solved by redesigning T7 expression systems to remove cryptic transcription. To do this, new sequence-to-function models are needed to predict where and how often T7 RNAP binds across a DNA template, coupled to a computational optimization algorithm that removes undesired T7 promoter sites hiding within other genetic elements, including inside protein coding sequences. In our prior work, we developed a sequence-to-function model that predicts transcriptional initiation rates for endogenous bacterial RNAP/σ^70^, called the Promoter Calculator, and demonstrated that this predictive design capability can be used to lower cryptic transcription in engineered genetic systems^27, 28^. However, a similar sequence-to-function model has not yet been developed for T7 RNAP. Prior engineering efforts have mostly focused on maximizing transcriptional initiation by designing new canonical T7 promoters^29, 30, 31, 32, 33^. To date, there is no comprehensive sequence-to-function model that quantifies T7 RNAP’s interactions with arbitrary DNA sequence and predicts its transcription rate from both canonical and low-affinity promoters.

Here, we present a sequence-to-function model, called the T7 Promoter Calculator, that accurately predicts the T7 RNAP transcriptional initiation rate for every potential start site across an input DNA sequence, including both canonical and low-affinity promoters. We carried out a massively parallel reporter assay to develop this model by building 11588 T7 expression systems with systematically mutated T7 promoter sequences and measuring their transcriptional initiation rates in pooled IVT reactions, leveraging molecular barcoding and deep sequencing (**Figure 1A**). Using this data, we trained and tested a linear machine learning model to calculate T7 RNAP’s binding free energy (ΔG_total_) to each potential transcriptional start site (**Figure 1B**). We combined the predictive model with computational optimization to redesign a T7 expression system that produces the coxsackie and adenovirus receptor (hCXADR) protein, removing undesired T7 RNAP binding sites during the design of the 5’ UTR and coding region (**Figure 1C**). We applied direct RNA nanopore sequencing to quantify the frequency of transcription starts and termination stops in IVT reactions, measuring RNA purity from natural versus redesigned expression systems. Overall, we found that redesigning expression systems using the T7 Promoter Calculator removed several low-affinity sites, resulting in lower cryptic transcription and higher RNA purity.

**Figure 1.**
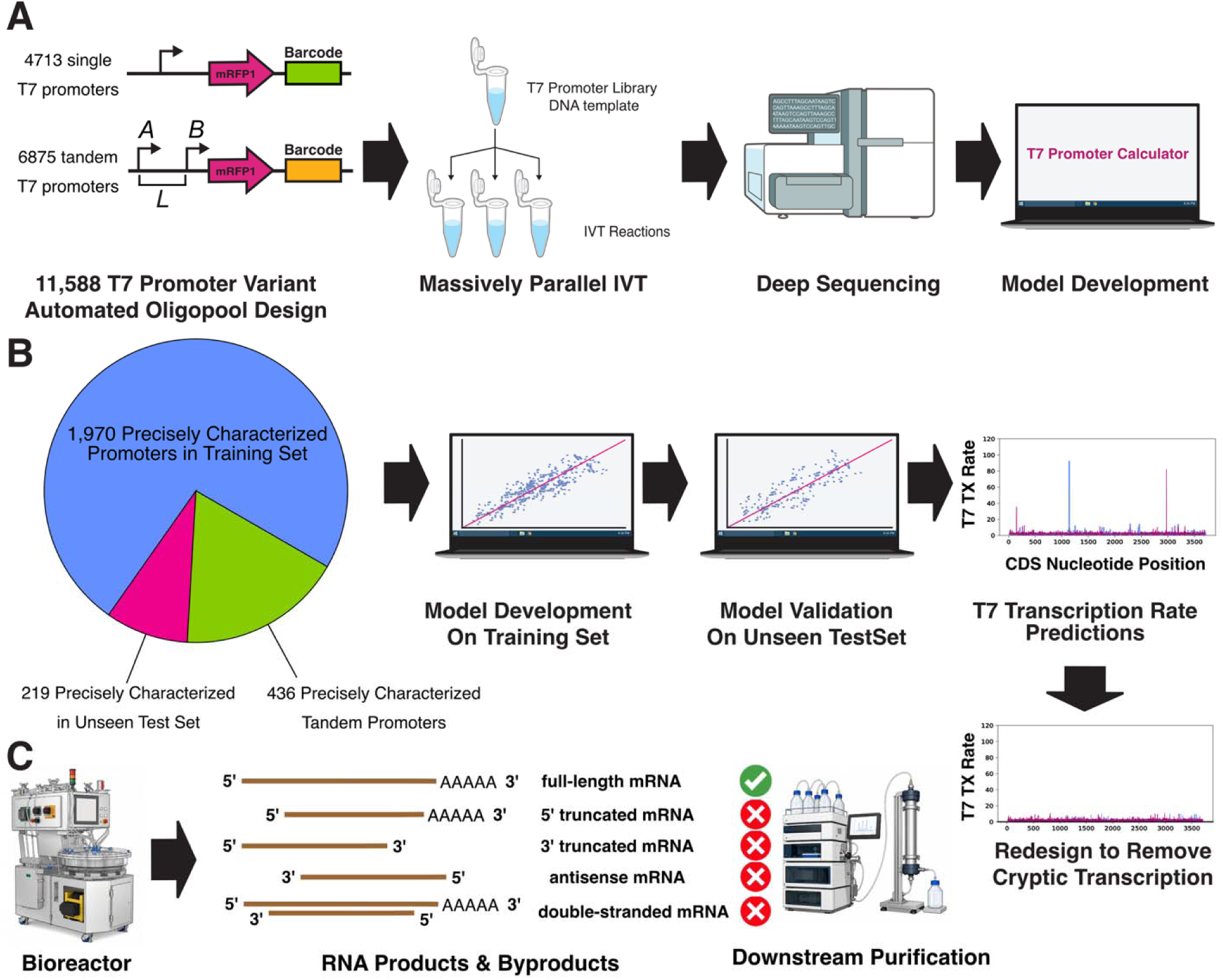
Development of a T7 Promoter Calculator model using a massively parallel design-build-test-learn workflow. **A** 11588 single and tandem T7 promoters were designed, built, and tested, combining *in vitro* transcription (IVT) reactions and deep sequencing for massively parallel measurements. **B** A T7 Promoter Calculator model was parameterized and validated on training and unseen test datasets. The model is combined with generative design to remove cryptic transcription from a T7 expression system. **C** Biomanufacturing of therapeutic RNAs produces many undesired RNA byproducts that are costly to remove during downstream purification. The T7 Promoter Calculator is used to redesign T7 expression systems to decrease cryptic transcription and improve the purity of full-length mRNAs.

## Results

### Design and construction of a library of T7 promoters and expression systems

The canonical T7 promoter is a 23 bp sequence that has highly conserved contacts to T7 RNAP starting from the -17 position and ending at the +6 position^34, 35, 36, 37, 38, 39^. Mutational studies and evolutionary analysis show that nucleotide substitutions in the binding and recruitment domain (- 17 to -6), particularly at -11G, -9C, -8T, and -7C, will significantly lower transcription rates, while mutations in the DNA melting region (-4 to -1) will reduce transcription rates under certain conditions^40, 41, 42, 43, 44, 45^. Once bound, T7 RNAP enters several rounds of productive and abortive cycling, generating short mRNA transcripts (10-12 nt) in an attempt to form a stable R-loop. After a stable R-loop is formed, T7 RNAP transitions from initiation to processive elongation^39, 42^. Prolonged rounds of cycling can reduce the T7 promoter’s transcription rate and increase the production of short RNA transcripts.

Additional sequence motifs outside the canonical promoter region will also affect transcription rates^29, 30, 35, 36, 41, 43^. For example, the first transcribed nucleotides have been shown to facilitate R-loop formation, including a +1 motif GGGAGA at the beginning of the mRNA transcript and the absence of pyrimidines from +1 to +8. Upstream AT-rich motifs positioned from -22 to -17 may increase the rate of T7 RNAP recruitment. Additionally, there is some evidence that the - 18C position is conserved and important to controlling transcription rate. Notably, both the core promoter sequence and the surrounding flanking regions are likely to play a critical role in controlling transcription rate, motivating a systematic investigation.

We therefore designed a library of T7 promoters that included: 4521 T7 promoter sequences with systematically mutated positions to measure positional effects; 6785 combinations of 2-promoter tandem designs to test transcriptional additivity, T7 RNAP saturation, and roadblocking; and 195 natural and previously engineered T7 promoters for comparisons. Starting from a baseline T7 promoter (T7Max)^29, 41^, we systematically introduced substitutions to create T7 promoter sequence variants: all possible 1-bp and 2-bp mutations within the 23 bp core region (-17 to +6), creating 2346 sequence variants; all possible 5-bp mutations within two 5-bp upstream regions (-27 to -23 and -22 to -18), creating 2048 sequence variants; and all possible 3-bp mutations within two 3-bp downstream regions (+7 to +9 and +10 to +12), creating 128 sequence variants (**Figure 2A**). We then selected pairs of canonical T7 promoters and placed them in same-direction tandem designs, separated by a neutral spacer sequence with a systematically varied length (0 to 100 bp), creating 6785 sequence variants. Finally, we added engineered T7 promoters optimized for high transcription, several promoter variants containing >2 consensus mutations or alternative flanking sequences, and the 17 natural T7 promoters from the T7 bacteriophage genome^16, 29, 30, 41, 43, 44, 45^. T7 promoter sequences with excluded cut sites were removed from the library designs. The overall library contains 11588 T7 promoter sequences with well-defined features. The complete sequence specifications are available in **Supplementary Data 1**.

**Figure 2.**
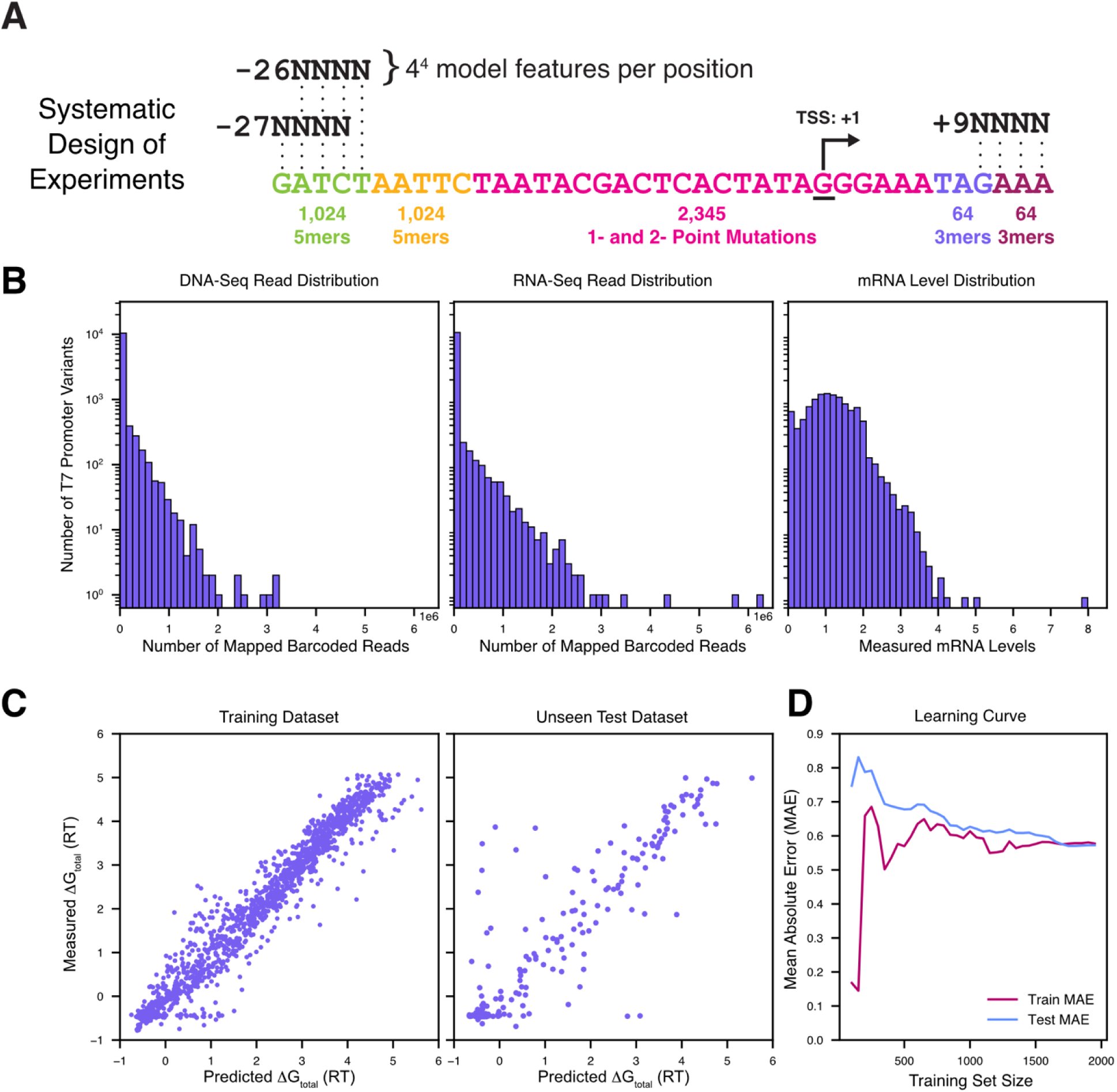
T7 Promoter Design, Measurements, and Model Performance. **A** Schematic shows the systematic design of single T7 promoter variants, varying nucleotide positions and k-mer motif sequences. **B** DNA-Seq and RNA-Seq measurements are used to quantify the mRNA levels across T7 promoter variants. **C** Training and validation of the T7 Promoter Calculator model, showing predictive accuracy across the training (Pearson R^2^ = 0.96, *p* < 10^-80^) and unseen test datasets (Pearson R^2^ = 0.80, *p* = 10^-78^). **D** Learning curves show the mean absolute error (MAE) when using a subset of the training dataset to parameterize the model, followed by validation on the unseen test dataset. Learning curves achieve convergence with similar MAEs (train 0.52 vs. test 0.56) before training data is exhausted.

To build the library, we carried out 2-step DNA assembly to create 11588 plasmids that each use a designed T7 promoter to express a protein coding sequence (mRFP1) with a unique barcode located in the 3’ untranslated region (**Methods**). Here, mRFP1 is not used as a reporter; it is only being used to create a full-length mRNA transcript. The barcode sequences were designed by the Oligopool Calculator to be pair-wise distinguishable with up to 3 mismatches^50^. The positioning of the barcode within the 3’ UTR prevents it from interfering with the transcription rate measurements. To build this library, an oligopool with 11588 oligonucleotides was synthesized and inserted into a vector (ColE1, CmR) using: (i) PCR to amplify the oligopool template and produce a DNA fragment library; (ii) restriction digestion with SacI and BmtI to cut the DNA fragment library at outer flanking sites; and (iii) ligation between the pre-digested vector and DNA fragment library. The vector contained a downstream T7 terminator to efficiently terminate transcription. We also confirmed that the vector does not have any sequence motifs resembling a T7 promoter. In the second DNA assembly step, a RBS-mRFP1 constant DNA fragment was inserted into the plasmid library in between the T7 promoter and the barcode, using: (i) restriction digestion with SalI and BamHI to cut the plasmid library at inner insertion sites; and (ii) ligation between the digested plasmid library and pre-digested RBS-mRFP1 fragment (**Methods, Supplementary Figure 1**). We then used long read nanopore sequencing to assess library construction; over 90% of mapped reads showed correct sequences.

### Massively parallel measurements of a library of T7 expression systems

We began by using DNA-Seq to measure the composition of the T7 expression system library. We carried out a low-cycle PCR on the plasmid library to generate barcoded amplicons, followed by short read sequencing (Illumina). We used the Oligopool Calculator to carry out read filtering and barcode mapping, returning over 8.8 million mapped reads^46^. We identified 10906 system variants (94% library coverage) with a broad distribution of DNA-Seq read counts. 2467 system variants had at least 10 DNA-Seq read counts, while 1209 system variants had at least 1000 DNA-Seq read counts (**Figure 2B**).

We then carried out IVT reactions using the T7 expression system library to produce a pool of barcoded mRNA transcripts, followed by RNA-Seq. Extracted RNA was converted to cDNA, followed by a low-cycle PCR to generate barcoded amplicon libraries and deep short read sequencing (Illumina). We applied the Oligopool Calculator to map over 4 billion reads, identifying between 500 million to 700 million mapped barcoded reads per sample with an overall library coverage of 11588 system variants and at least 30 RNA-Seq read counts per variant. We quantified the mRNA levels for each system variant by dividing the RNA-Seq read counts by the DNA-Seq read counts across the triplicate datasets (**Figure 2B**). We then applied a stringent filter on this dataset to remove measurement noise, only including system variants if they had at least 1000 DNA-Seq read counts. Overall, this dataset contained 1209 T7 expression system variants with measured mRNA levels (N = 3 separate IVT reactions as replicates) that varied over a 6300-fold dynamic range.

We then analyzed an additional 677 characterized T7 promoter variants, measured from previous sequencing efforts^35, 45^, that contained combinations of sequence mutations that produced lower-affinity T7 promoters. Overall, this supplementary T7 promoter dataset had transcription rates that varied across a 4500-fold dynamic range with an average transcription rate that was 22-fold lower than our dataset. Importantly, both datasets contained transcription rate measurements of the same natural T7 promoters, enabling us to use these as reference measurements and providing a straight-forward way to carry out cross-dataset normalization. We therefore normalized the transcription rate measurements across both datasets, creating an augmented dataset containing 1828 T7 expression system variants with transcription rates that varied across a 14000-fold dynamic range. The sequences, deep sequencing measurements, and normalized transcription rates for all T7 expression system variants are included in **Supplementary Data 2**.

### Training and testing a sequence-to-function model to predict T7 transcription rates

We first formulate a relationship between the measured read-out (mRNA level) and an intrinsic biophysical property of the T7 promoter sequences (Gibbs binding free energy, ΔG_total_). In our IVT reactions, there is a pool of T7 RNA polymerase competitively binding to a plasmid library containing a pool of T7 promoter sequences. Through competitive binding, a T7 promoter sequence with a more negative ΔG_total_ will bind to T7 RNAP with a higher probability, according to the Boltzmann relationship of equilibrium statistical thermodynamics, which states that the probability of binding is proportional to exp (-*β*ΔG_Total_), where *β* is the Boltzmann constant for a system with constant temperature and pressure (here *β =*1.636 *mol/kcal*). Using this relationship, the measured transcripts’ mRNA levels (m_i_) can be converted to the T7 promoters’ apparent binding free energies (ΔG_total,i_) according to ΔG_Total_,_i_ log (m_i_) /*β*.

Our IVT reactions were designed to satisfy the equilibrium criteria, including: (i) the amount of T7 RNAP was limiting, which largely prevents promoter saturation; (ii) the probability that T7 RNAP binds to any T7 promoter is determined by the T7 promoter’s sequence and controls its recruitment rate; (iii) the IVT reaction was run long enough to carry out many cycles of transcriptional initiation and elongation so that the number of mRNAs produced is proportional to the initiation rate without significant accumulation of intermediate states; and (iv) there are no RNases or ribosomes in the IVT reaction that would alter mRNA levels.

To train and test the sequence-to-function model, we randomly split the augmented dataset into 1609 sequence variants in the training set and 219 sequence variants in the unseen test set. We formulated linear machine learning models (ridge regression) that use different sequence-encoded features to predict the apparent ΔG_total_ of each T7 promoter sequence. For each feature set, we trained the ridge regression model’s coefficients using the training dataset and then evaluated its predictions on the unseen test set. We also calculated learning curves where the training step is carried out with limiting amounts of datapoints, followed by evaluation on the test set, comparing their mean absolute errors (MAE). We then systematically varied sequence-encoded feature sets and carried out hyperparameter optimization until the MAE learning curve on the training dataset converged to the MAE learning curve on the unseen test set (**Supplementary Figure 2, Supplementary Data 3**). This procedure ensures that the machine learning model learns a compact, generalized relationship that limits the number of trainable parameters and prevents memorization.

We found that position-indexed, overlapping 4-mers across the 39-bp promoter sequence window yielded the most accurate and generalized model. Each nucleotide in a 36-bp window is encoded as a 4-bp motif (left-positioned). The presence or absence of each 4-bp motif at each position is a categorical (binary) feature in the model. There are 64 possible 4-bp motifs each at position, returning a linear model with 2304 trainable parameters; however, because the 4-bp motif sequences are overlapping, there are functional constraints between these parameters that must be satisfied during training. Using this featurization, we found that the model accurately predicted the apparent ΔG_total_ of the T7 promoter sequences in the unseen test set (Pearson R^2^ = 0.80, two-tailed T-test *p* = 10^-78^) (**Figure 2C**). The learning curves achieved convergence with a MAE of 0.52 for the training set and a MAE of 0.56 for the unseen test set, showing that there was sufficient data to parameterize this model with low error (**Figure 2D**).

When applied to the unseen test set, the model (called the T7 Promoter Calculator) accurately predicted T7 transcription rates with an average error of 2-fold across a large 500-fold range. The model error was uniform across this range, covering both high-affinity and low-affinity T7 promoters. For example, there were 92 T7 promoter variants in the test set with higher transcription rates than the baseline T7Max (ΔG_total_ = -1 to 0); for these promoters, the model’s average fold-change error was only 1.47. At the other extreme, there were 54 T7 promoters with very low affinity (20 to 50-fold lower transcription rates than T7Max); importantly, the model was still able to predict the transcription rates of these low-affinity T7 promoters with an average fold-change error of 2.6. Overall, 73% of the T7 promoter sequences were well-predicted with a fold-change error of 2 or less (**Supplementary Data 2**).

We then analyzed the T7 Promoter Calculator model to quantify the maximum and minimum change in ΔG_total_ (model-predicted ΔΔG_total_) when mutating each nucleotide in the 36-bp sequence window (-27 to +9). As expected, we found that the most sensitive nucleotide positions were found within the core binding and DNA melting regions (-17 to +1) (**Figure 3A**). The most sensitive positions are -11, -9, -8, and -7, which is consistent with prior work^17^. We then analyzed the top 20 and bottom 20 4-mer motifs, ranked by their ΔΔG_total_ (**Figure 3B**). For example, we found that the highest affinity 4-mer motifs were TCAC starting at position -8 and ACTC starting at position -10. Together, they show that a core TC motif at position -8 has a potent effect on T7 RNAP binding affinity with minor contributions from flanking nucleotides. The model also identified 4-mer motifs that greatly penalized T7 RNAP binding affinity, including

**Figure 3.**
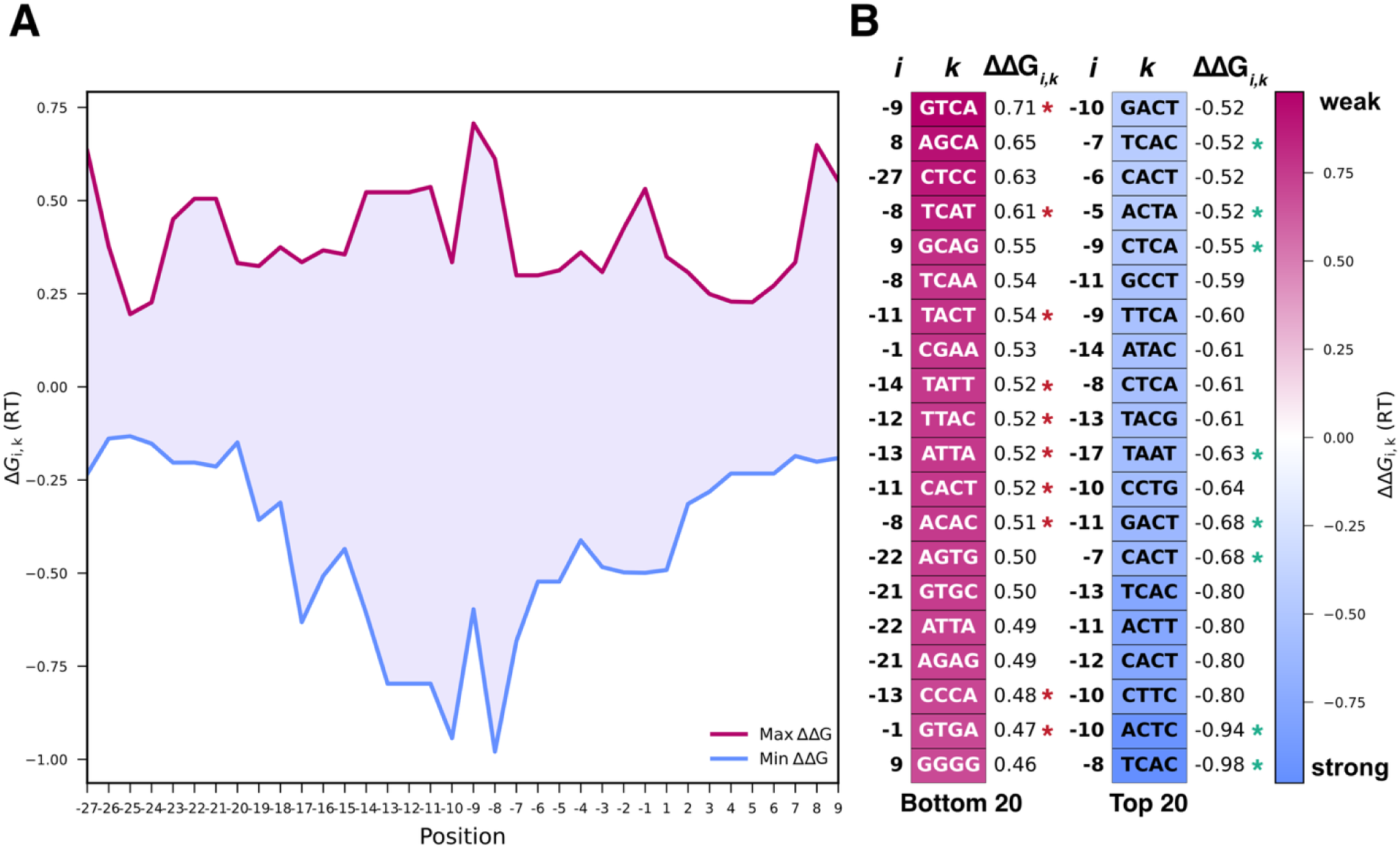
Model analysis reveals motif sequences controlling T7 transcription rates. **A** The model-predicted minimum and maximum binding free energy (ΔΔG_total_) contributions at each sequence position, indexed with respect to the predicted transcriptional start site (+1). **B** The rank-sorted motif sequences controlling T7 transcription rates, showing their start sites, motif 4-mer sequence, and color-coded binding free energy contributions. Green asterisks denote known, conserved nucleotide positions. Red asterisks denote positions with known deactivating mutations.

TACT starting at position -11 (containing a deactivating -11T), a GTCA starting at position -9 (containing a deactivating -9G), and ACAC starting at position -8 (containing a deactivating -8A). Notably, there are several important 4-mer motifs outside the core binding region, including the AT-rich motif TAAT at position -17 that increases transcription rate, consistent with prior work^35^. Thus, the T7 Promoter Calculator model recapitulates known canonical interactions, while expanding our knowledge to lower-affinity, non-canonical motifs located inside the core region as well as in the outer flanking regions.

### Model-predictive design removes cryptic T7 transcription and improves mRNA purity

We next demonstrate how the T7 Promoter Calculator model can improve the manufacturing of therapeutic RNAs by removing cryptic transcription and increasing the purity of full-length mRNA. We combined the T7 Promoter Calculator model with a generative design algorithm to scan across a genetic system sequence, identify undesired T7 promoter sites in both the forward and reverse directions, and introduce nucleotide changes to minimize the T7 transcription rate to below a very low threshold (**Methods**). When nucleotide changes are part of a protein coding sequence, only synonymous codons are introduced. Additional design rules can be applied to ensure high expression levels of the therapeutic RNA inside human primary cells.

As a first application of this algorithm, we redesigned a T7 therapeutic mRNA expression system that encodes human coxsackie and adenovirus receptor (hCXADR or hCAR) protein. hCXADR is the receptor protein used by adenovirus to enter cells, though many tissues (including solid tumors) do not express enough CXADR to enable adenovirus-mediated gene delivery, limiting the uptake and effectiveness. By first introducing the therapeutic mRNA into these tissues (as LNPs), transient expression of CXADR can promote adenoviral uptake and gene therapy effectiveness. The T7 Promoter Calculator predicts that the natural CXADR coding sequence contains 20 low-affinity T7 promoter sites across both strands; 8 in the forward (sense) direction and 12 in the reverse (antisense) direction (**Figure 4A**). The generative design algorithm introduced over 900 synonymous nucleotide changes to the CXADR coding sequence to remove 19 low-affinity T7 promoter sites, prioritizing the removal of antisense sites that produce dsRNA and sense sites that produce polyA-tailed RNA products (**Figure 4BC, Supplementary Data 4**). The target threshold for removing cryptic transcription was to limit T7 RNAP binding at any potential site to ΔG_total_ > 9. We also identified and removed a T7 RNAP pause motif (5’-ATCTGTT-3’) that promotes premature transcriptional termination. We synthesized and constructed these T7 expression systems (**Methods**).

**Figure 4.**
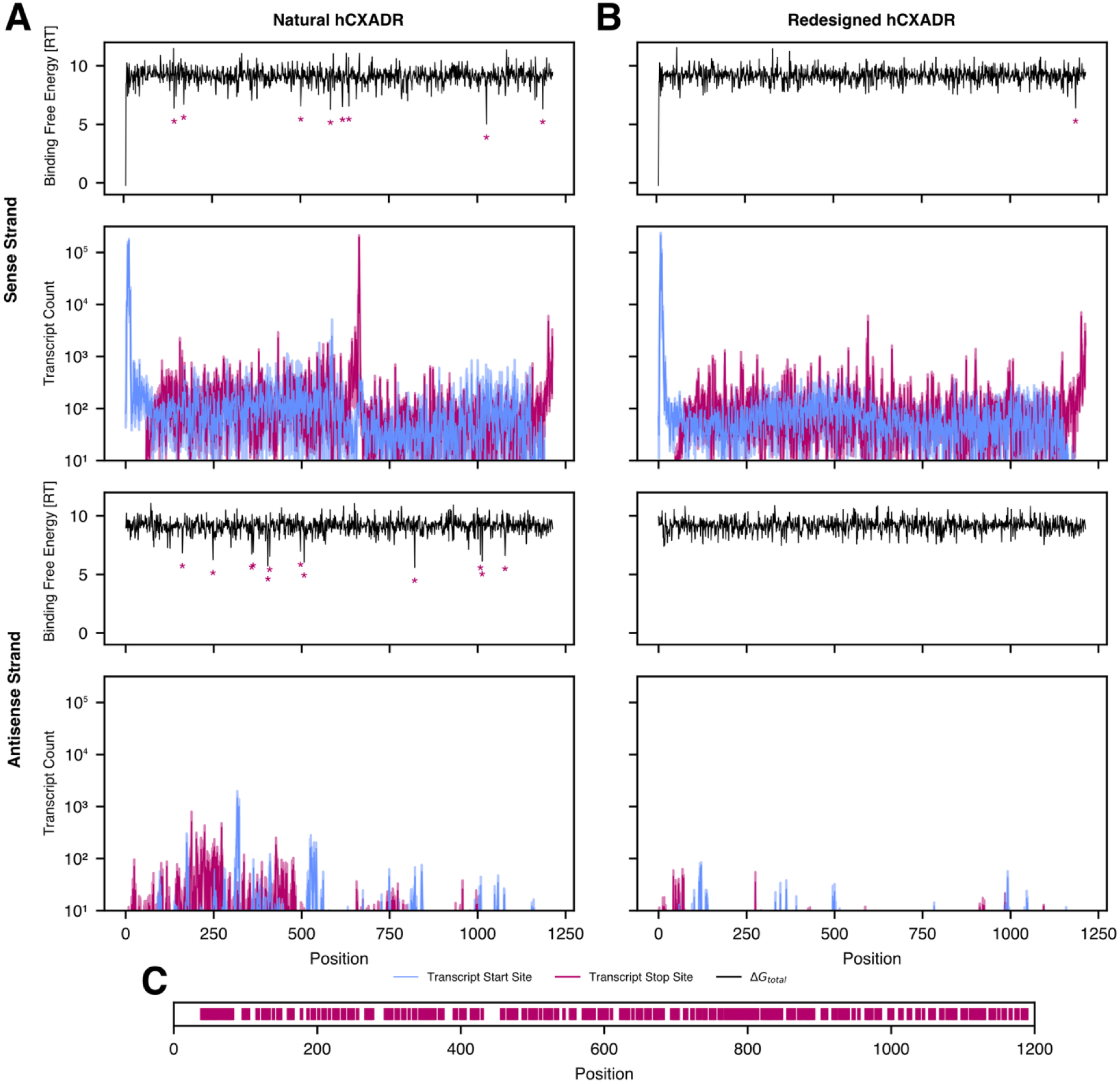
Redesign of a T7 expression system to remove cryptic transcription. **A** The T7 Promoter Calculator is used to predict T7 RNAP binding free energies (ΔG_total_) across the (top) sense and (bottom) antisense strands of a T7 system expressing the natural hCXADR protein. Red asterisks (*) show predicted low-affinity transcriptional start sites. The measured frequencies of (blue) transcriptional start sites and (red) transcriptional stop sites are shown below the predictions. Lines are the mean of biological replicates across each nucleotide position. Shaded regions show the 68% confidence intervals. **B** The T7 Promoter Calculator was used to carry out generative design of the hCXADR nucleotide coding sequence to remove cryptic T7 RNAP binding sites (ΔG_total_ < 9 RT), introducing over 900 synonymous nucleotide changes. Model predictions and site-specific transcriptional frequency measurements show a large reduction in RNA byproducts, including a 1.65-fold higher titer of full-length mRNA (*p* = 0.042), a 2.1-fold higher purity of full-length mRNA (*p* = 0.003), and a 14.5-fold lower amount of free antisense RNA (*p* = 0.03). **C** Nucleotide substitutions (red lines) were introduced to redesign the CXADR CDS sequence with lower model-predicted T7 transcription rates. The redesigned CXADR shares 74% sequence similarity with the natural CXADR sequence.

We then carried out identical IVT reactions to produce the natural and redesigned hCXADR mRNAs (N = 3 replicates x 2 DNA templates), using the same canonical T7 promoter for transcription of the full-length mRNA. To measure where T7 transcription actually starts and stops across the DNA templates, we utilized MinION nanopore sequencing with the RNA flow cell and Direct RNA sequencing kit (**Methods**). Direct RNA sequencing carries out sequencing and base calling of single RNA strands without the need to convert to cDNA or carry out fragmentation. We then map each read sequence, extract transcriptional start and stop sites, and count the number of transcription initiation and termination events taking place across each nucleotide position (**Figure 4AB**). Notably, Direct RNA sequencing does not produce reads from double-stranded RNA; any reads from antisense RNAs were captured in their single-stranded state.

As expected, for both systems, Direct RNA sequencing shows a high transcriptional initiation rate in the sense direction at the canonical T7 promoter position with a positional offset of about 10 nt. However, when applied to the natural hCXADR sequence, Direct RNA sequencing revealed a substantial amount of cryptic transcription in both the sense and antisense directions with a clear termination signal located nearby the T7 RNAP pause site (**Figure 4A**). Notably, downstream sense start sites will produce truncated RNA byproducts that will contain polyA tails if they are not prematurely terminated, while downstream antisense start sites will produce antisense RNA that will form double-stranded RNA product. In contrast, Direct RNA sequencing showed that the redesigned hCXADR sequence produced significantly less truncated and antisense RNA at downstream sites (**Figure 4B**). Across both sequences, the production of antisense RNA is overall lower than sense RNA as a large fraction of the antisense RNA is likely in double-stranded form.

We calculated the mean and standard deviation of four metrics to quantify and compare these measurements: (i) the number of full-length mRNA transcripts (titer); (ii) the fraction of full-length mRNA over all RNA produced (purity); (iii) the fraction of nucleotide positions with ultralow transcription (positional purity); and (iv) the fraction of single-stranded antisense RNA over full-length mRNA (antisense impurity). The redesigned hCXADR sequence produced full-length mRNA with a 1.65-fold higher titer (3.3 ± 0.5 x 10^5^ vs 2.0 ± 0.3 x 10^5^, *p* = 0.042) and a 2.1-fold higher purity (58 ± 0.04% vs 27 ± 0.05%, *p* = 0.003) as compared the natural hCXADR sequence. The positional purity of the redesigned hCXADR sequence was increased by 1.67-fold (70 ± 0.03% vs 42 ± 0.05%, *p* = 0.004) and its antisense impurities were greatly reduced by 14.5-fold (0.4% ± 0.1% vs 5.8% ± 1.7%, *p* = 0.03). Overall, the redesigned hCXADR sequence has a significantly higher full-length purity with a much lower antisense RNA proportion. The removal of the T7 RNAP pause site greatly reduced premature termination, increasing full-length mRNA titer.

### Designed single and tandem T7 promoters testing maximal transcription and rate-limiting steps

Across our entire dataset, we characterized 660 T7 promoter variants that had higher transcription rates than the baseline T7Max promoter. In particular, the T7 promoter “V-1280” with sequence 5’-GATCTGGGGATAATACGACTCACTATAGGGAAATAGAAA-3’ contained mutations at positions -22 and -18 (outside the core binding region) that resulted in a 3.6-fold higher transcription rate than T7Max. The T7 Promoter Calculator model predicted its binding free energy as ΔG_total,predicted_ = -0.62 as compared to its apparent ΔG_total,measured_ = -0.78, which is within the mean model error. Here, we designate this promoter as T7Skramz.

In parallel, we wondered whether 2-promoter tandem designs would increase T7 transcription rates beyond a single promoter. Tandem designs will also test whether adjacent promoter regions could independently recruit T7 RNAP and carry out transcriptional initiation in an additive fashion or whether the promoter regions would become saturated with bound T7 RNAP, limiting the maximum possible transcription rate. Excess DNA supercoiling from nearby promoters could also limit maximal transcription rates. We designed and characterized 76 tandem 2-promoter sequences, selecting pairs of previously tested T7 promoters with high transcription rates, and combining them together in the same orientation, separated by a spacer region. Across the pairs of T7 promoters, we systematically varied the length of the spacer from 0 bp (no spacer) to 100 bp in order to quantify the effect of steric hindrance on independent T7 RNAP binding.

Interestingly, we found that 2-promoter tandem designs generally had lower measured transcription rates than either of the two individual T7 promoters (**Supplementary Figure 3**). There were about 17 tandem designs that had very low transcription rates (more than 2-fold lower than T7Max), even though their individual promoters were measured to have high activities. Excluding these outliers, there is still a sub-additive relationship when comparing the sum of the individual T7 promoters’ measured transcription rates to the 2-promoter tandem designs’ measured transcription rates with a proportionality constant of about 0.20 (Pearson R^2^ = 0.33, *p* = 2x10^-6^, **Supplementary Data 2**). We tested whether the transcription rate of the upstream T7 promoter or the downstream T7 promoter had a greater effect on the tandem 2-promoter region; we found that both promoters contributed about equally to the 2-promoter region’s overall transcription rate with proportionality constants of 0.23 and 0.17, respectively. Instead, we found that the minimum of the two promoters’ individual transcription rates had the best ability to explain the transcription rate of the tandem promoter region (Pearson R^2^ = 0.43, *p* = 2x10^-8^) with a proportionality constant of about 0.40. Interestingly, we did not observe any effect from changing the spacer length. Overall, this analysis suggests that placing two high-activity T7 promoters in tandem creates a rate-limiting transcriptional bottleneck where the lower-activity promoter controls the tandem promoters’ cumulative transcription rate. Even beyond this rate-limiting step, there is an additional penalty for placing two T7 promoters close to each other (tested up to 100 bp apart).

Two separate mechanisms could be responsible for the observed penalties in transcriptional activity. First, high-activity T7 promoters generate a significant number of positive DNA supercoils in the forward direction and negative DNA supercoils in the reverse direction, creating a promoter positional dependency. Increasing the upstream T7 promoter’s transcription rate could potentially repress the transcription rate of the downstream T7 promoter by increasing its DNA supercoiling density, increasing the amount of work to form a stable R-loop. Conversely, increasing the downstream T7 promoter’s transcription rate would have a beneficial effect on the upstream promoter’s transcription rate. Notably, inside IVT reactions, there are no gyrases or topoisomerases to alter DNA supercoiling densities or resolve topological knots. Alternatively, if transcriptional initiation was rate-limited by a blocking intermediate state, for example, DNA melting and the closed-to-open conformational change or during R-loop formation, then the presence of this blocking intermediate state in either promoter could create a rate-limiting step for both promoters in a tandem orientation. Overall, in this dataset, we do not see any promoter positional effect, but we do see observe a shared, rate-limiting step controlled by their minimum transcription rate. Thus, placing two T7 promoters in tandem will not increase their overall transcription rate as they will likely share the same rate-limiting step during transcriptional initiation.

## DISCUSSION

We developed a sequence-to-function model, called the T7 Promoter Calculator, that predicts where and how often T7 RNA polymerase initiates transcription across a DNA template inside *in vitro* transcription reactions. To do this, we designed, built, and tested 11588 T7 promoters, combining massively parallel IVT reactions, molecular barcoding, and deep sequencing to measure their mRNA levels. Overall, these T7 promoters varied transcription rate by a 6300-fold dynamic range with systematic perturbation of the interactions that control recruitment, DNA melting, and R-loop formation. Using this data, we trained and validated a machine learning model with high accuracy and generalizability (Pearson R^2^ = 0.80 on the unseen test dataset, two-tailed T-test *p* = 10^-78^). The model predicts the T7 transcription rate at each potential start site across a long DNA sequence, including both canonical T7 promoter sequences and low-affinity T7 binding sites that contribute to cryptic transcription. The model considers a longer 39-bp sequence region, including flanking motif sequences, that were found to additionally control T7 transcription rates. The T7 Promoter Calculator can be used to predict, design, control, and optimize genetic system sequences to achieve target T7 transcription rates at the desired transcriptional start sites, while assessing and redesigning to ensure that all other potential start sites have minimally low T7 transcription rates.

As a demonstration, we applied the T7 Promoter Calculator to decrease cryptic T7 transcription during the production of a therapeutic mRNA encoding the human coxsackie and adenovirus receptor (hCXADR) protein. Redesign of this expression cassette increased full-length mRNA purity by 2.1-fold and reduced the detectable production of antisense RNA by 14.5-fold in IVT reactions, utilizing Direct RNA nanopore sequencing to count each transcriptional start and stop site. Lowering the production of RNA byproducts is important as truncated and anti-sense RNAs are a key source of adverse immunogenicity inside the human body. By greatly reducing the production of these RNA byproducts, one can streamline the design, manufacturing, and purification of mRNA therapeutics, reducing the number of purification steps, decreasing batch times, reducing downstream costs, and mitigating potential adverse effects. Importantly, the predominant cost of manufacturing therapeutic RNAs is their downstream purification^26^. These improvements are synergistic within existing RNA therapeutic developmental platforms, where it is possible to rapidly proceed from a target protein to a clinically ready formulation.

The model-based design of T7 expression systems can incorporate multiple rules and objectives, in addition to removing cryptic T7 transcription, to achieve desired functions and outcomes. For example, eukaryotic 5’ UTR sequences have been designed to maximize translation initiation rates by incorporating canonical Kozak motif sequences, removing upstream open reading frames (uORFs), and decreasing the formation of inhibitory mRNA structures^47, 48^, while prokaryotic 5’ UTR sequences are well-designed by the RBS Calculator model to control translation rates^49^. Eukaryotic 3’ UTR sequences have been designed to maximize mRNA stability and accelerate the formation of circular mRNAs by adding RNA-binding protein (RBP) motif sequences, including polyA binding protein (PABP)^50^, while prokaryotic 3’ UTR and intergenic sequences are well-designed by biophysical models that calculate end-dependent and end-independent mRNA decay rates^51, 52^. Multi-objective design algorithms have been created to generate genetic system sequences that satisfy multiple criteria^28^, for example, ensuring that therapeutic RNAs are well-expressed inside human cells after delivery, while also ensuring that their manufacturing achieves high mRNA titers and full-length purity.

Past efforts at reducing the production of RNA byproducts have focused on improved T7 RNA polymerases and optimizing the IVT reaction itself. For example, the C-terminal domain of T7 RNA polymerase was computationally redesigned to accelerate the transition from transcriptional initiation to elongation^53^, which produces fewer short RNAs during R-loop formation. In another example, thermostable T7 RNA polymerases were used within high-temperature IVT reactions to reduce the formation of double-stranded RNA and 3’ extension products^54^. The IVT reaction process and composition have also been extensively optimized to maximize mRNA titer and purity, including the introduction of denaturants^55, 56, 57^. Enzyme and process improvements can be synergistic, although the coupling between upstream and downstream processes can also introduce trade-offs. For example, a higher temperature IVT reaction can lead to mRNA degradation, while the addition of denaturants can cause precipitants to form inside preparative columns. Thus, the ability to lower the production of long RNA byproducts by removing cryptic transcription provides an additional and synergistic approach to improving full-length mRNA purity.

Interestingly, prior efforts have not investigated how the presence of low-affinity T7 sites within DNA templates could have contributed to the observed production of long RNA byproducts. The number of low-affinity T7 sites within a DNA template will vary across applications, introducing a previously unexplained source of process variability during biomanufacturing. Here, the T7 Promoter Calculator can be used to predict *a priori* when and where a DNA template will bind to T7 RNAP at undesired sites, enabling researchers to redesign those sites and eliminate a key source of RNA byproducts before any experiments are conducted. Notably, engineered T7 RNA polymerases often retain the same DNA binding domain as natural T7 RNAP and thus bind to DNA sites with the same model-predicted binding free energies.

T7 RNA polymerase is also widely used as an orthogonal transcriptional resource within engineered genetic circuits across different organisms^33, 58, 59^. T7 promoters are engineered to activate transcription in response to environmental signals or to dynamically control metabolic processes^31, 32^. The T7 Promoter Calculator can be used to predict the effects of flanking sequences on a T7 promoter’s transcription rate, design synthetic T7 promoters with targeted transcription rates to tune functions, and determine where there are low-affinity T7 sites inside the genetic circuit that would unexpectedly break the circuit’s function. The widespread use of the T7 Promoter Calculator model promises to further improve the designability and reliability of engineered genetic circuits that are powered by T7 RNAP.

A Python source code implementation of the T7 Promoter Calculator is available at https://github.com/hsalis/SalisLabCode. A web interface to the T7 Promoter Calculator is available at https://salislab.net/software.

## METHODS

### T7 promoter library and oligopool design

We designed an 11588 T7 promoter library beginning with a 39 bp T7 promoter sequence (V-1), containing the consensus T7 promoter adjoined by constant upstream and downstream sequences to serve as the background sequence of all T7 promoter variants. A custom Python code was written to generate 4519 T7 promoter variants consisting of: (1) 2345 1- and 2-point mutations in the 23 bp sequence at positions [-17, +3] relative to the TSS, (2) 1024 5-mers at position -27, (3) 1024 5-mers at position -22, (4) 64 3-mers at position +7, and (5) 64 3-mers at position +10. An additional 195 T7 promoters were sourced from the literature, including all wild-type T7 promoters in the bacteriophage T7 genome, additional T7 promoter sequences with 3 or more mutations, and T7 promoter sequences with varied upstream and downstream sequences. 6875 tandem T7 promoters were designed with varied promoters, positioning, and spacer lengths (0 to 100 bp, increments of 10 bp).

Oligopool design was automated using the Oligopool Calculator to design barcodes, primer binding sites, and padding sequences. Each oligonucleotide contains: (1) a T7 promoter variant, (2) SacI, SalI, BamHI, and BmtI digest cutsites, (3) an 11 bp barcode sequence, (4) 2 primer binding sites, (5) a 6 bp spacer sequence for mRFP1 insertion, and (6) a padding sequence to equalize each oligo to a length of 250 nt. The oligopool was ordered from Twist Biosciences. A list of all T7 promoter and oligo sequences can be found in **Supplementary Data 1**.

### T7 promoter library cloning

The pFTV1 vector modified (pFTV1-O1) with a landing pad cassette containing a T7 terminator, SacI and BmtI restriction digest cutsites, and a reverse primer binding site sequence for PCR amplification of sequence barcodes was used for oligopool cloning. The pFTV1-O1 vector was double digested with SacI and BmtI restriction enzymes (New England Biolabs, NEB) and purified via gel electrophoresis. The oligopool was PCR amplified with Primer 1 and Primer 2 for 14 cycles and double digested with SacI and BmtI and purified via Zymo DNA Clean and Concentrator Kit (Zymo Research). The oligopool was ligated into pFTV1-O1 to create pFTV1-O2. pFTV1-O2 was digested with SalI and BamHI restriction enzymes and purified via gel electrophoresis. The mRFP1 was PCR amplified from the pFTV1 vector with primers containing SalI and BamHI primer arms, respectively, and treated with DpnI (NEB) followed by gel electrophoresis purification. The mRFP1 amplicon was digested with SalI and BamHI and ligated into the linearized pFTV1-O2 vector. The resulting product was a plasmid library containing the T7 promoter variants upstream of the mRFP1 sequence and a unique barcode. The plasmid library was then transformed into in-house prepared DH10B electrocompetent cells, recovered for 1 hour in SOC, transferred to selective LB media + 20ug/mL chloramphenicol, and cryostocked after overnight growth. All PCR and ligation reactions were completed with Q5 High-Fidelity DNA Polymerase (NEB) and T7 DNA Ligase (NEB). Annotated sequences may be found **Supplementary Data 1**.

### Plasmid library *in vitro* transcription measurements

The plasmid library was linearized via NotI-HF digestion (NEB) and purified via gel electrophoresis and retained in a single aliquot. DNA-Seq quantification of the T7 promoter variants was carried out by PCR amplifying the barcode region of the linearized plasmid library with 10ng of DNA template for 14 cycles using Primers 5 and 6.

Triplicate *in vitro* transcription reactions were carried out using 120ng of linearized plasmid library template with HiScribe T7 High Yield RNA Synthesis Kit (NEB) for 2 h. Reactions were treated with TURBO DNase Kit (Invitrogen) to remove DNA template and purified using RNA Clean and Concentrator Kit (Zymo Research). To measure RNA levels of T7 variants, RNA-Seq measurements were executed by first-strand cDNA synthesis using LunaScript RT Master Mix Kit (NEB) with Primer 2. Barcoded regions were PCR amplified from 10ng of the reverse transcribed product using Primer 2 and Primer 3 for 14 cycles. Sequencing was completed via custom amplicon sequencing of 150 bp paired-end reads (GeneWiz) for RNA-Seq and DNA-Seq measurements. Barcodes were mapped and counted using the Oligopool Calculator to obtain RNA-Seq and DNA-Seq read counts for T7 promoter variants. The measured mRNA levels are the RNA-Seq read count of each replicate divided by the DNA-Seq read count of the input library. The error in the mRNA levels is the standard deviation of the 3 biological replicates.

### Building a ridge regression model for predicting T7 transcription rates

At thermodynamic equilibrium, the probability of T7 RNA polymerase binding to a T7 promoter within a pool of plasmids follows a Boltzmann distribution:

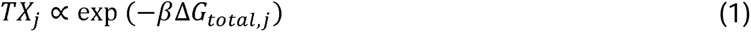

where ΔG_total,j_ is the Gibbs binding free energy of T7 RNA polymerase bound to the j^th^ T7 promoter, *β* is the Boltzmann constant, and TX_j_ is the transcription rate of the j^th^ T7 promoter. By selecting a reference promoter with a reference transcription rate measurement (TX_ref_) and binding free energy (ΔG_total,ref_), we can convert the proportional relationship into a definitive one:

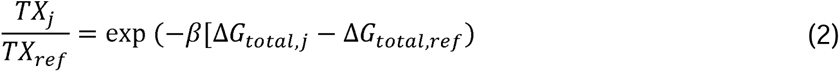

We selected V-11544 (T7Max) as the reference promoter to create an ideal comparator. We arbitrarily define ΔG_total,ref_ = 0 as the reference point to simplify an equation to calculate the apparent Gibbs binding free energy of the j^th^ T7 promoter:

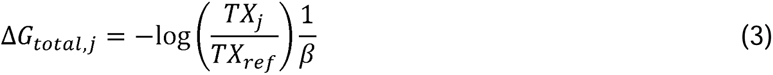

We use sliding motif features across the DNA sequence query. During model training, each motif at each position is assigned a coefficient, which is its additive contribution to ΔG_total,j_. The ΔG_total,j_ of a DNA sequence is then computed using:

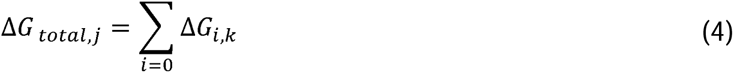

where *i* is the nt position of the DNA query and *k* is the k-mer motif appearing at position *i*.

To train the model, we first create a stringent filter that excludes any measurements with fewer than 1000 DNA-Seq reads. We then split the filtered dataset into a training set (90%) and unseen test set (10%). We trained a ridge regression model with Python (v3.12.4) using Scikit Learn’s RidgeCV function (v1.5.1). The 39 bp T7 promoter variant sequences were one-hot encoded as overlapping sequential 4-mers to generate model feature inputs. Outcomes were the measured ΔG_total,j_ values. We carried out 5-fold cross validation, repeated 10 times, to determine the optimal regularization parameter (α). Following model training, we identified outliers in the training set (datapoints with >3 MAE). We then repeated model training on the training set, ignoring these outliers. The final model was used to predict ΔG_total,j_ on the unseen test dataset. Statistical comparisons and metrics were carried out using Scikit-Learn.

### Identification and removal of cryptic T7 transcription in the CXADR gene sequence

hCXADR sequence was accessed from UniProt (P78310) and the National Center for Biotechnology Information (NCBI EMBL: BT019876). A custom Python code was written to implement an iterative scan-and-replace search, applying our model to predict ΔG_total_ at each position in the hCXADR gene and substitute synonymous codons in-frame within the sequence query when the predicted ΔG_total_ was too favorable (less than 9 RT). This procedure continued until all sites had unfavorable binding free energies, resulting in a redesigned hCXADR sequence. DNA fragments were synthesized (Integrated DNA Technologies) containing the T7 ϕ6.5 promoter, a 5’ UTR sequence, and either the natural hCXADR or redesigned hCXADR sequences. DNA fragments were inserted into the pFTV1-O1 vector, as previously described. Plasmids containing the natural and redesigned hCXADR sequences were linearized via NotI-HF digestion (NEB) and purified via gel electrophoresis. Triplicate IVT reactions were carried out on 120ng of linearized template using HiScribe T7 High Yield RNA Synthesis Kit (NEB) for 1 hour at 0.5x T7 RNAP concentration. RNA was purified as previously described. 10 ug RNA was polyadenylated using *E. coli* poly(A) polymerase (NEB) for 1.5 minutes at 37°C. Polyadenylated RNA was purified with Agencourt RNAClean XP beads (Beckman Coulter) and sequenced with Oxford Nanopore Direct RNA Sequencing Kit (SQK-RNA004) (Oxford Nanopore Technologies) on a FLO-MIN004RA flow cell with a MinION Mk1D. Sequencing was controlled by MinKNOW software (v24.11.10) until estimated sequenced bp reached 0.75 Mbp. A minimum Q score threshold of 10 was used to remove low-quality reads. Custom Python code was used to align the first 100 nt and last 100 nt of each read to reference RNA sequences, followed by mapping the transcriptional start sites and transcriptional termination stop sites. Measured transcriptional start sites were found to be right-shifted by 10 nt as compared to model-predicted start sites. A uniform 10-nt left-shift was applied to the measured transcriptional start sites.

## DATA AVAILABILITY

All promoter sequences, model weights & calculations, experimental measurements, and supplementary statistical analysis are available in Supplementary Data 1, 2, 3, and 4.

## CODE AVAILABILITY

We used Python v3.12.4 and custom code that employed the following Python modules to develop the T7 Promoter Calculator: sklearn v1.5.1, SciPy 1.16.3, numpy v2.3.5, pandas v2.3.3, and matplotlib v3.10.3. A Python source code implementation of the T7 Promoter Calculator is available at https://github.com/hsalis/SalisLabCode with Git release tag name r2026.

## ACKNOWLEDGEMENTS

This project was supported by funds from the National Science Foundation [DBI-2400302, MCB-2131923].

